# GTEx_Pro: A robust and accurate preprocessing pipeline for GTEx data: TMM+CPM normalization and SVA batch correction for enhanced multi-tissue analysis

**DOI:** 10.1101/2025.04.26.650748

**Authors:** D. Jothi

## Abstract

GTEx_Pro is a Nextflow-based pipeline for preprocessing GTEx v8 transcriptomic data, enhancing multi-tissue comparability. It integrates TMM + CPM normalization and SVA batch effect correction to improve biological signal recovery while reducing systematic variations across 54 GTEx tissues. Designed for scalability and reproducibility, GTEx_Pro facilitates accurate multi-tissue transcriptomic analysis and a similar framework can be adapted to other large-scale transcriptome datasets.

## Main

Integrating and analyzing large-scale biological datasets is crucial for understanding complex biological processes and disease mechanisms. Public resources such as Genotype-Tissue Expression (GTEx) provide extensive gene expression data across diverse human tissues, enabling studies on gene expression trajectories, tissue-specific gene correlations, and expression quantitative trait loci (eQTL) [1, 2]. However, the cross-sectional nature of the GTEx dataset, where the tissue samples are derived from different individuals of the age group 20-79 of varying locations, introduces technical artifacts and batch effects based on donor demographics and tissue/sample processing. GTEx_Pro, a preprocessing pipeline (Fig 1), addresses these challenges by integrating Trimmed Mean of M component (TMM) normalization, Counts Per Million (CPM) scaling, and Surrogate Variable Analysis (SVA) batch correction [3, 4]. TMM removes library size differences and compositional biases, while CPM allows gene expression comparisons across samples by scaling the adjusted counts. SVA further mitigates batch effects, improving the reliability of downstream gene expression analysis. Our results demonstrate enhanced biological signal recovery and reduced technical artifacts compared to other processing methods. Leveraging Nextflow, GTEx_Pro ensures automation, scalability, and reproducibility across computational environments. Its open-source design facilitates applications in diverse transcriptomic studies. Hence, GTEx_Pro supports more accurate multi-tissue gene expression analysis and downstream research applications by improving data cleaning.

**Figure 1:**
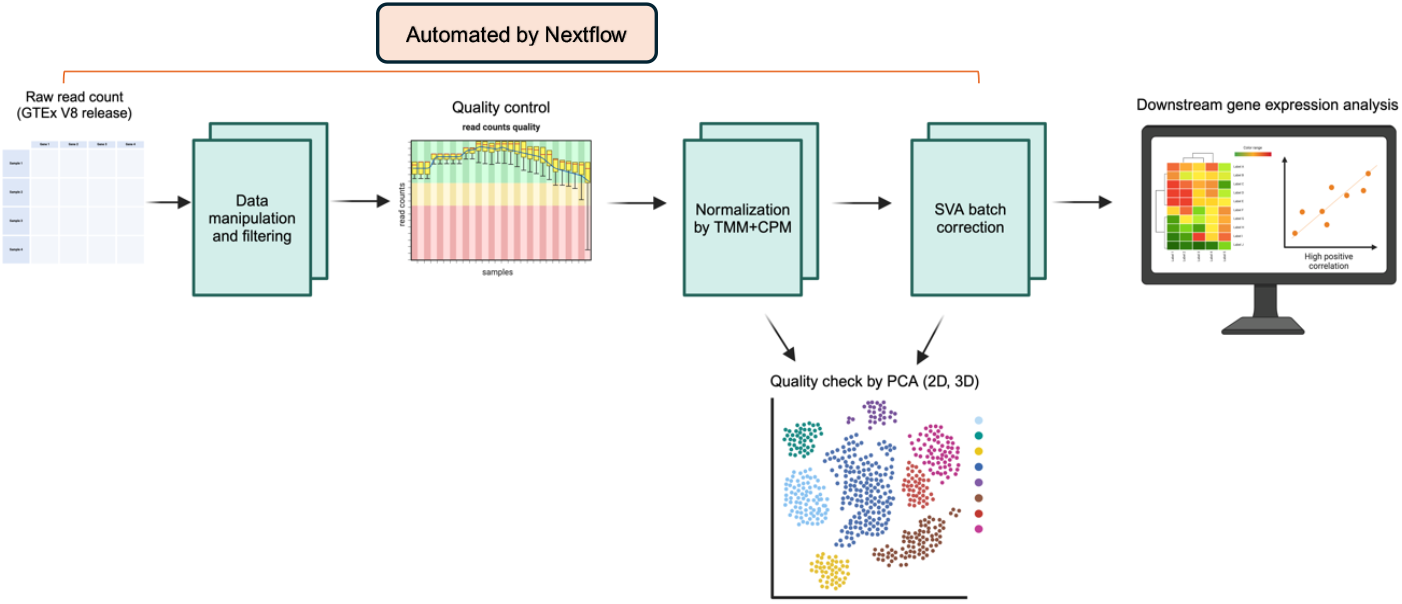
GTEx_Pro Preprocessing Workflow for Downstream Analysis of GTEx Tissues. This diagram illustrates the preprocessing steps performed in GTEx_Pro to analyze GTEx tissue datasets. The workflow begins with downloading the raw read count matrix, which undergoes data manipulation and filtering to pre-select a specific set of genes for further processing. Furthermore, quality control is performed by removing missing, invalid values (Inf/NaN) and any potential outliers manually, followed by normalization using the TMM and CPM methods. Batch effects are then addressed using SVA with sex as a covariate. PCA is employed to explore principal component variance and tissue clustering quality. The pre-processed data can be subsequently used for various downstream analyses, such as tissue-specific expression studies.

3D Principal Component Analysis (PCA) of the raw count data revealed substantial overlap between samples from different tissues, with clustering primarily driven by technical variation rather than biological signal (Fig. 2A). Following preprocessing by the TMM+CPM pipeline, 3D PCA showed a pronounced enhancement in tissue-specific clustering (Fig. 2A). Samples from the same tissue, such as Heart Atrial Appendages, Heart Left Ventricle, and Skin-SunExposed, Skin-NotSunExposed, grouped, reflecting improvements in the biological signal. Notably, biologically distinct tissues such as the brain, the Liver, the Heart, and the Skin were clustered separately as expected. However, the PC1 variance in TMM-normalized CPM values (TMM+CPM) was lower than that of raw read counts. This reduction is likely due to the scaling down of highly expressed genes during normalization, which adjusts for differences in sequencing depth and gene expression distribution. Normalization can occasionally lower principal component variance by minimizing the influence of highly expressed genes, thereby enhancing biologically relevant signals and improving tissue-based separation [5]. Furthermore, following SVA batch correction, the grouping of distinct tissue clusters was further apart than by TMM+CPM processing alone. This is evident from the increased Euclidean distance [6] between the tissue clusters, as shown in Fig. 2A. More specifically, the average Euclidean distance score between tissue clusters increased after SVA batch correction across all 54 GTEx tissues (Fig. 2B, Supplementary Fig. 1), suggesting the effectiveness of the batch correction method. In addition, the overall variance in the first two principal components is increased by 1.5% (Fig. 2C).

**Figure 2:**
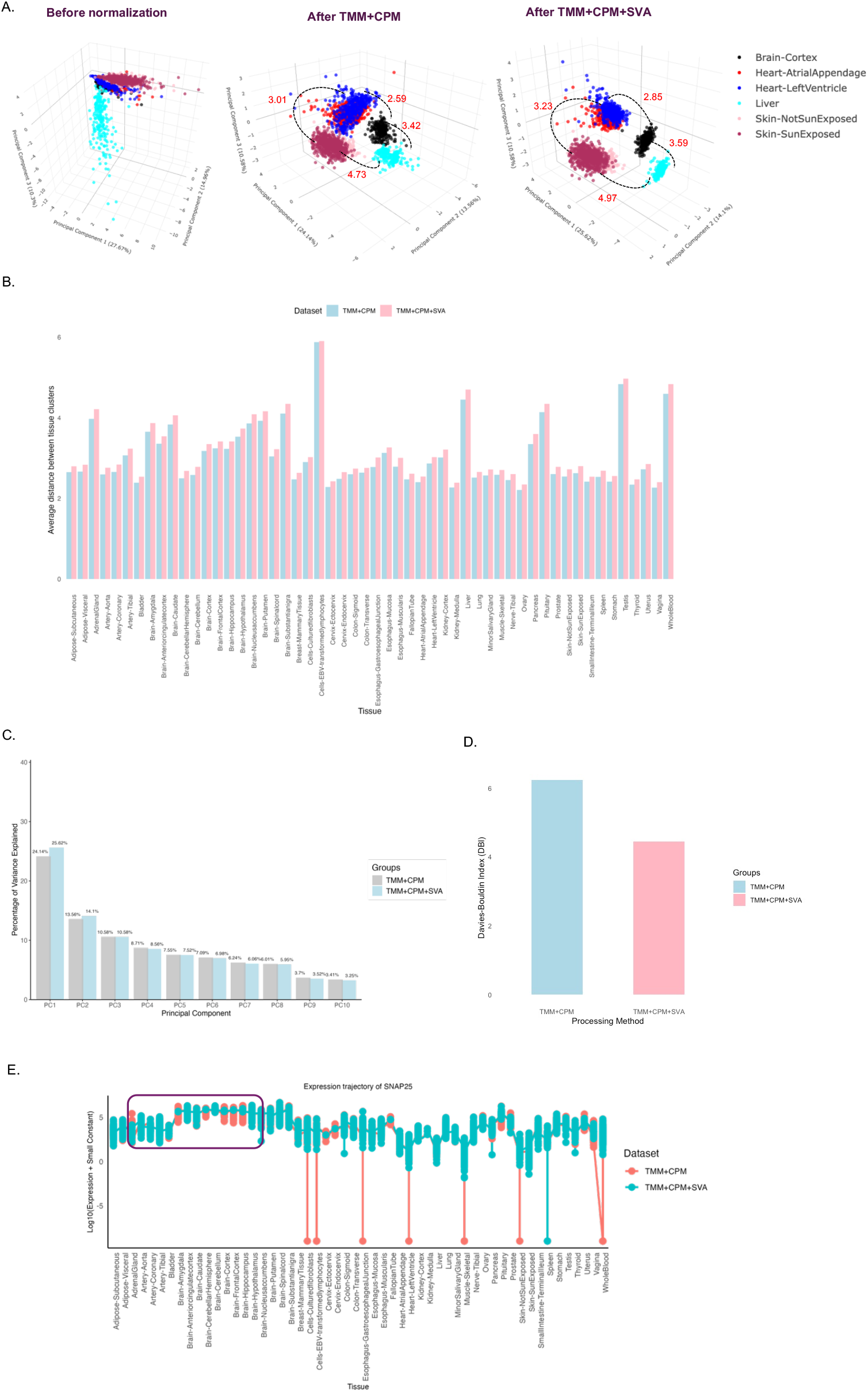
Comparison of clustering quality of the pre-processing pipeline TMM+CPM+SVA. A) 3D PCA plot displaying the separation of samples by tissue type before normalization, after normalization, and after batch correction. The tissue clusters of Brain-Cortex, Heart-Atrial Appendages, Heart-Left Ventricle, Liver, Skin-Sun Exposed, and Skin-Not Sun Exposed were shown as a representation and the measure of Euclidean distance between PCA tissue clusters is denoted in red text. The 3D plots can be visualized using the links (https://dhana2403.github.io/3D_plots/3d_pca_plot_all_tissues_tmm.html), (https://dhana2403.github.io/3D_plots/3d_pca_plot_all_tissues_sva.html). B) The bar graph compares the average Euclidean distance after TMM+CPM and TMM+CPM+SVA processing steps across 54 GTEx tissues. The centroid for each tissue is calculated as a mean of PC1, PC2, and PC3. C) The bar graph illustrates the percentage of variance explained after applying TMM+CPM normalization and SVA batch correction on gene expression data. The percentage of variance corresponding to the principal components in the x-axis is mentioned at the top of each bar. D) The bar graph compares the Davies-Bouldin Index (DBI) for tissue clusters after TMM+CPM normalization and SVA batch correction. DBI is calculated based on the first three principal components (PC1, PC2, and PC3). Lower DBI values indicate better separation and compact clusters. E) The plot compares brain tissue-specific gene SNAP25 expression trajectory after TMM+CPM and TMM+CPM+SVA processing steps across 54 tissue types. The distribution of expression values in brain tissue samples is highlighted in the purple box.

Furthermore, the tissue clustering quality is estimated by the Davies-Bouldin index (DBI) [7, 8] estimates the tissue clustering quality. DBI measures the average similarity ratio of each tissue cluster with its most similar cluster, where a lower DBI indicates better clustering. It was observed that after SVA batch correction, the DBI index score decreased, suggesting better clustering following the batch correction method (Fig 2D). Additionally, a trajectory plot of the brain-tissue-specific gene *SNAP25* was generated using the implemented workflow to assess the pipeline’s efficacy for downstream analysis. It was observed that the distribution of normalized value for brain tissue samples became more stable and consistent following the SVA batch correction method (Fig 2E). This pattern was also observed in other tissue-specific genes, such as albumin (ALB) in the Liver, Pancreas, and Whole Blood, and keratin (KRT1) in Skin-SunExposed and Skin-NotSunExposed tissues (Supplementary Fig 3). Moreover, these subtle improvements in PCA clustering quality lead to substantial alterations in the gene-gene correlation heatmap analysis (Supplementary Fig 5). Collectively, these findings suggest that even marginal enhancements in Euclidean distance and PCA clustering quality can substantially impact the reliability and interpretability of downstream gene expression analyses.

For benchmarking, traditionally used GTEx normalization methods, such as TPM, were compared to the TMM+CPM processing method, along with SVA batch correction or quantile normalization [9]. The comparison demonstrated that SVA batch correction plays a critical role in enhancing the pipeline’s effectiveness. Notably, the TMM+CPM+SVA and TPM+SVA approach outperformed TPM+quantile normalization and TMM+CPM+quantile in enhancing the Euclidean distance between PCA tissue clusters (Supplementary Fig 4). To further assess the impact of the GTEx_Pro preprocessing pipeline on downstream gene expression analysis, gene expression trajectories across 54 GTEx tissues were evaluated for all four processing methods. The results indicated that both TMM+CPM+SVA and TPM+SVA displayed stable and consistent gene expression values compared to other groups. However, gene expression values derived from the TMM+CPM+SVA pipeline were consistently higher than those from TPM+SVA, suggesting that TMM+CPM normalization enhances expression estimates while maintaining distributional consistency (Supplementary Fig 3 B). Additionally, the pipelines were cross-validated using an alternative set of genes, including AKT1, GSK3B, GDF11, FOXO1, SESN2, ULK1, PGC, PINK1, PDPK1, BCL2, HMOX1, FIS1, TNF, and PARP1. The results revealed that, upon SVA batch correction, the pipeline consistently improved the Euclidean distances between tissue clusters for most tissues, mirroring the findings from the previous analysis. This demonstrates the robustness and effectiveness of the pipeline in enhancing tissue-specific expression profiles for most of the GTEx data (Supplementary Fig 4).

Although the current pipeline is specifically designed for GTEx data, the underlying framework of TMM+CPM+SVA can be applied broadly to other RNA-seq preprocessing workflows. We anticipate that this approach will enhance downstream gene expression analysis and facilitate accurate tissue-specific drug discovery efforts [10]. Future pipeline developments will include integrating machine learning-based batch correction [11] methods to assess their effectiveness in further preserving biological signals. Additionally, a web-based platform to enable researchers to interactively visualize results generated using the GTEx_Pro preprocessing pipeline will be made available. A key limitation of the current pipeline is that it applies only to the GTEx dataset. Future updates will focus on addressing this limitation to expand the applicability of the framework for all types of tissue-specific analysis.

Overall, the pipeline effectively integrates existing normalization and batch correction methodologies while highlighting critical yet previously overlooked clustering variations that substantially impact downstream gene expression analysis. By addressing these factors, the pipeline improves the accuracy and reliability of gene expression analysis, thereby enhancing the robustness of subsequent biological interpretations

## Online Methods

### Data Acquisition and Processing

The GTEx project is a public resource providing gene expression data across a diverse range of human tissues. In this study, we utilized GTEx v8 RNA sequencing data, comprising 54 tissues with a total of 17,235 samples. The dataset includes RNA-seq data from individuals of varying ethnic backgrounds and sexes. The GTEx v8 data can be accessed through the dbGaP database (*https://gtexportal.org*) and is available in the form of raw read counts as well as normalized gene expression metrics (e.g., TPM, FPKM). To facilitate automated data retrieval, the GTEx_Pro pipeline directly downloads the raw read count file (GTEx_Analysis_2017-06-05_v8_RNASeQCv1.1.9_gene_reads.gct) along with associated metadata files(GTEx_Analysis_v8_Annotations_SampleAttributesDS.txt) and (GTEx_Analysis_v8_ Annotations_SubjectPhenotypesDS.txt) from the GTEx portal.

### Metadata Processing

Following data acquisition, sample metadata was processed in R using the data: table, tidyverse, and dplyr libraries. Key metadata attributes were extracted and cleaned, including sample ID, tissue type, batch effects, RNA integrity number (RIN), and ischemic time. Subject phenotype data was processed separately, with categorical variables converted into appropriate factor levels. The metadata files were merged based on the subject ID, and unique records were retained after filtering.

### Gene Expression Data Processing

Raw gene expression counts were imported using fread(), and gene identifiers were standardized by removing version numbers. The dataset was filtered to retain only the genes of interest as defined by the user. Infinite values were replaced with NA, and rows containing missing data were excluded. Additionally, columns (samples) exhibiting zero variance were discarded to eliminate low-information data.

### Tissue-Specific Filtering and Data Storage

Samples were grouped by tissue type, ensuring that each group contained a minimum of two replicates. The processed expression data for each GTEx tissue was stored in .rds format in the directory data/processed/expression/readcounts_all/. The metadata was saved separately as attphe_all.rds for further downstream analyses.

### Data Normalization by TMM

To normalize gene expression data, we applied the TMM normalization method using the edgeR package. TMM normalization corrects for differences in sequencing depth and composition biases across samples, ensuring comparability of gene expression levels. Each tissue-specific read count file is loaded along with its corresponding metadata. Samples flagged as outliers (if specified) are excluded. The dataset is then filtered to retain only samples present in both the metadata and read count matrices. For each tissue, the normalization factors were calculated using calcNormFactors(), which applies TMM normalization [12]. Furthermore, normalized values were scaled using the CPM () function. The final normalized values were saved in .rds format in the output directory. The summary table containing the sample counts for each tissue was compiled and saved as a CSV file (sample_counts.csv) (Supplementary Table 1).

### Quantile normalization

To ensure the comparability of expression values across samples and mitigate technical variation, quantile normalization was applied to transcript-level expression matrices on a per-tissue basis. The input data consisted of pre-normalized matrices (e.g., TPM or TMM-normalized CPM) derived from RNA-seq datasets. Both sample metadata and expression matrices were loaded into R, and outlier samples identified a priori, were excluded. Only samples present in both the expression and metadata files were retained. Quantile normalization was performed using the normalize.quantiles() function from the preprocessCore package [13]. This method aligns the empirical distribution of expression values across samples by sorting, averaging ranks, and reassigning values such that all columns share an identical distribution. Following normalization, the gene and sample identifiers were restored, and the normalized matrices were saved in .rds format for downstream analyses. The number of samples retained per tissue after filtering was recorded. In this study, quantile normalization was benchmarked with two distinct normalization pipelines: 1) TMM + CPM + Quantile Normalization and 2) TPM + Quantile Normalization. Although quantile normalization is more commonly applied alongside TPM to ensure comparability of expression values across samples, we also assessed its performance when combined with TMM-normalized CPM values to evaluate whether it provides consistent results in mitigating technical variation and improving the comparability of gene expression levels across different RNA-seq normalization strategies.

### Batch Correction by SVA

To correct for batch effects and hidden confounders in gene expression data, we employed Surrogate Variable Analysis (SVA) and the removeBatchEffect() function. The number of surrogate variables (SVs) was estimated using the num.sv() function with the ‘be’ method [14], a technique based on parallel analysis. In the case of sex-specific tissues, SVA was omitted as the model incorporates sex as a covariate, with batch effect correction performed using removeBatchEffect(), where batch1 was included as a covariate. Tissues with fewer than 20 samples were excluded from batch effect correction due to the insufficient sample size for reliable adjustment. The batch-adjusted gene expression data for all tissues were saved in .rds format within the adjusted_sva_all directory.

### Principal Component Analysis (PCA)

#### Interactive 3D Plot Visualization

Normalized gene expression data were obtained from multiple tissues, each represented as an individual file. Negative values, if present, were replaced with a small positive constant (1e-9) to ensure numerical stability. Data matrices from all tissues were then merged into a single dataset. PCA was performed using the prcomp() function in R [15], with centering and scaling applied to standardize gene expression values across tissues. The variance explained by each principal component (PC) was computed and reported. A three-dimensional PCA plot was generated using plotly [16] to visualize sample clustering across tissues. Principal components 1, 2, and 3 were selected, and the variance explained by each was annotated accordingly. Specific colors were assigned to predefined key tissues (e.g., *Brain-Cortex, Heart-AtrialAppendage, Liver*) for representation, while colors for remaining tissues were randomly assigned for distinct visualization. The interactive PCA plot was saved as an HTML file.

#### Quantification of Tissue Similarity Using Cluster Distance Metrics

To assess tissue similarity and the effect of batch correction on gene expression variation, we computed the Euclidean distances between tissue clusters in principal component (PC) space, using both uncorrected (TMM+CPM) and batch-corrected datasets (TMM+CPM+SVA). Tissue cluster centroids were computed as the mean coordinates of PC1, PC2, and PC3 for each tissue. Pairwise Euclidean distances between tissue centroids were calculated to quantify tissue similarity for TMM+CPM and TMM+CPM+SVA approaches. The distance matrices were converted into the long format and merged for comparative visualization. The average inter-tissue distance was computed for each tissue across both datasets. A bar plot was generated using ggplot2 to compare the average distance between tissue clusters under the two approaches. These distances demonstrate how “separate” the tissues are from each other in the PCA space. For instance, if tissues A and B have centroids that are very close to each other in PCA space, the Euclidean distance between A and B will be small, indicating that the two tissues are more similar in terms of the principal components. If tissues A and C are farther apart in PCA space, the Euclidean distance will be larger, indicating greater dissimilarity.

#### Variance Computation

For each approach, normalized values were log-transformed (with negative values replaced by a small positive constant, 1e-9). PCA was then performed using the prcomp() function in R, with both centering and scaling enabled. The percentage of variance explained by each principal component (PC) was computed from the eigenvalues of the covariance matrix as:

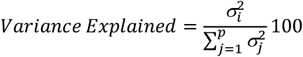

where 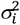 represents the variance of the ith principal component, and the denominator corresponds to the sum of all PC variances. The variance explained by the first 10 PCs was extracted for visualization.

#### Clustering Quality Assessment Using the Davies-Bouldin Index

To evaluate the impact of batch correction on clustering quality, we computed the Davies-Bouldin Index (DBI) for transcriptomic data processed with two different approaches. For each approach, normalized values were log-transformed, and PCA was performed using prcomp(). The first three principal components (PC1–PC3) were extracted for clustering assessment. Clustering quality was evaluated using the Davies-Bouldin Index (DBI), a measure of intra-cluster similarity and inter-cluster separation. The DBI was computed as:

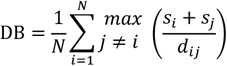

Where N is the number of clusters, *s*_*i*_ and *s*_*j*_ represents within cluster scatter (average distance of points to their cluster centroid) and *d*_*ij*_ denotes the inter-cluster distance between centroids. Lower DBI values indicate better clustering quality, implying greater separation between tissue clusters. Clusters were assigned based on tissue labels, and DBI values were computed using the index.DB () function from the clusterSim R package [17].

### Gene Expression Trajectory Analysis

To investigate the impact of the batch correction on downstream analysis, we analyzed the expression of tissue-specific genes like albumin (ALB), keratin (KRT1), and Synaptosomal-associated protein 25 (SNAP25)(Fig 2E, Supplementary Fig 2) in both batch-corrected and uncorrected datasets. For each approach, tissue-specific .rds files were loaded, and the expression values of the gene of interest were extracted and log10-transformed. To compare the impact of batch correction on gene expression values across multiple tissues, a line plot was generated using ggplot2 [18]. The tissue types were used as categorical variables on the x-axis, while the log-transformed gene expression values were plotted on the y-axis. Lines connect mean expression values per tissue for each normalization method. Points represent individual tissue-specific expression values for both TMM+CPM and TMM+CPM+SVA processing methods. The results demonstrate a direct comparison of gene expression trends across tissues of processing methods TMM+CPM and TMM+CPM+SVA, highlighting potential shifts introduced by SVA correction in tissue-specific genes.

### Correlation Analysis

Gene-gene correlation analysis was performed on tissue-specific gene expression datasets processed using the TMM + CPM and TMM + CPM + SVA pipelines. For each tissue, gene expression data was filtered to remove genes with zero variance. Spearman correlation was computed between gene pairs using the cor() function [19] with the “spearman” method and “pairwise.complete.obs” to handle missing values. Missing correlation values were replaced with zero. Heatmaps were generated with hierarchical clustering of genes and samples using the pheatmap package [20]. The resulting heatmaps were saved in the respective directory.

### Pipeline Implementation by Nextflow

To automate the three core modules, we utilized Nextflow 24.10.3 (https://www.nextflow.io), an open-source workflow manager widely adopted in bioinformatics [21], to construct a scalable and reproducible pipeline for transcriptomic data preprocessing. Nextflow serves as the backbone of the pipeline, organizing the execution of discrete tasks and managing data flow between them. The core components of Nextflow, including processes, workflows, and executors, enable the smooth orchestration and execution of the pipeline. Each step of the pipeline is encapsulated within its Nextflow process, ensuring modularity and a clear definition of system resource requirements.

The data acquisition process handles the extraction of raw gene expression data and associated metadata, based on a predefined list of genes of interest given by the user. This process is executed by an R script that retrieves and stores the data in a specified output directory. The second step in the pipeline, data normalization, applies TMM+CPM normalization to the raw gene expression data. In this process, an R script performs the normalization, using the previously acquired raw expression data. The output from this step includes the normalized gene expression values and sample count data. The final step in the pipeline is SVA batch correction, which removes unwanted sources of variation in the gene expression data due to batch effects. This process ensures that the data used for downstream analysis is adjusted for technical variation that could otherwise confound the results. As with the normalization process, SVA batch correction is implemented through an R script. The input consists of the normalized expression data, and the output is the adjusted gene expression values, ready for further analysis.

The Nextflow workflow component integrates these processes into a coherent pipeline, managing the flow of data between them. The workflow begins with the data acquisition process, followed by data normalization, and concludes with SVA batch correction. This data flow ensures that each step receives the necessary inputs and generates the required outputs, which are passed seamlessly to the next step. Once the pipeline is defined, Nextflow’s executor component manages the execution across the computational infrastructure. The workflow can be executed across various platforms, including local machines, cloud environments, or HPC clusters.

To run the pipeline, users simply need to navigate to the directory containing the workflow.nf file and execute the following command in the bash:

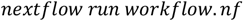

#### Pipeline Executions

The pipeline’s overall execution time was approximately 3 minutes and 48 seconds, with a total CPU usage of 0.1 CPU hours. This highlights the pipeline’s computational efficiency for evaluating a subset of high-dimensional data and performing complex preprocessing tasks such as data acquisition, normalization, and batch correction.

### Validation & Benchmarking

#### Cross-Validation

Cross-validation techniques were employed to assess the generalizability and robustness of the preprocessing pipeline. To ensure independent validation, a distinct set of genes (AKT1, GSK3B, GDF11, FOXO1, SESN2, ULK1, PGC1, PINK1, PDPK1, BCL2, HMOX1, FIS1, TNF, and PARP1), which were not initially used, was introduced. The resulting data was subsequently visualized using PCA, and Euclidean distance between tissue clusters was compared (Supplementary Fig. 4) to examine the consistency of the pipeline’s performance across new gene sets.

#### Benchmarking

Benchmarking was performed by obtaining TPM values from the GTEx dataset, which were then processed to eliminate technical variation either by SVA batch correction or quantile normalization. The Euclidean distance between tissue clusters was calculated and compared across different preprocessing groups to evaluate clustering quality. Additionally, tissue-specific expression trajectories were assessed across multiple preprocessing strategies, including: 1) TMM + CPM + SVA, 2) TPM + SVA, 3) TMM + CPM + Quantile Normalization, and 4) TPM + Quantile Normalization, to determine the impact of normalization and batch effect correction methods on clustering and trajectory patterns.

## Acknowledgments

I want to thank Friedrich Schiller University Jena for enabling open-access funding for my project.

## Authors contribution

DJ: conceptualization, coding, and manuscript writing.

## Code availability

The R code for the preprocessing pipeline and guidelines for executing the Nextflow workflow are available at https://github.com/dhana2403/GTEx_sample.

## Data availability

GTEx data can be accessed directly from https://gtexportal.org/. The data used for the analysis described in this manuscript were obtained from the GTEx Portal (https://gtexportal.org/) on 23/08/24.

