## Supplemental files for "GTEx_Pro: A robust and accurate preprocessing pipeline for GTEx data: TMM+CPM normalization and SVA batch correction for enhanced multi-tissue analysis"

Supplementary Figures

Fig S1:

A.

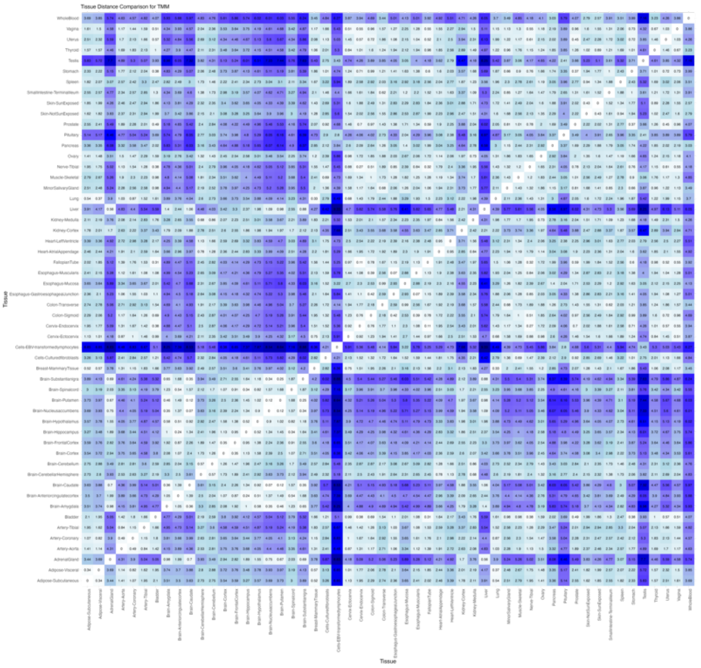


B.

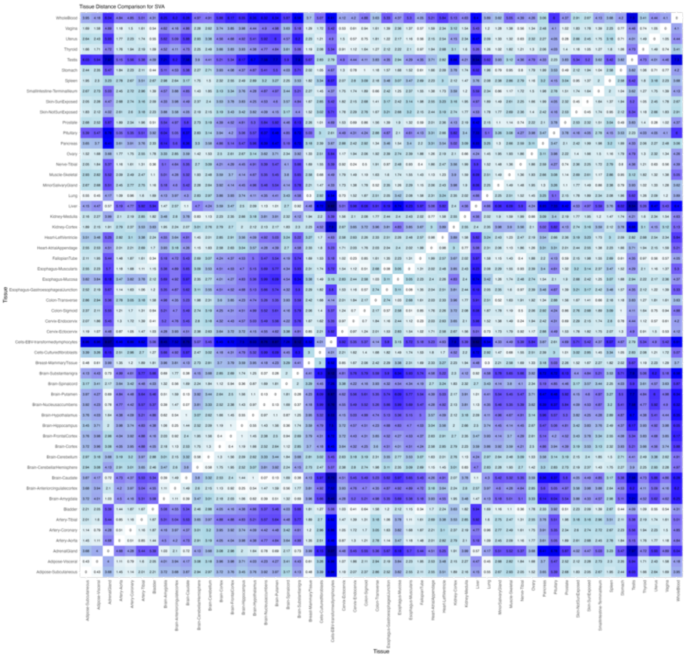


**Fig S1**: **Heatmap of pairwise Euclidean distance scores across 54 GTEx tissue clusters.** A) **After Normalization by TMM+CPM** B) **After TMM+CPM+SVA**. The heatmap scale represents Euclidean distance between tissue clusters, with darker shades indicating greater distance and dissimilarity and lighter shades indicating higher similarity and short distance between PCA tissue clusters.

Fig S2:

A.

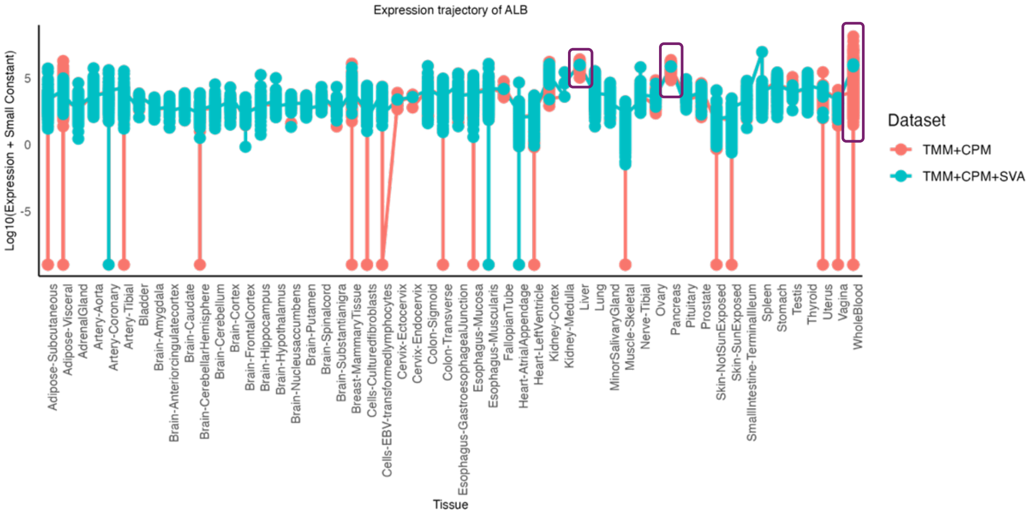


B.

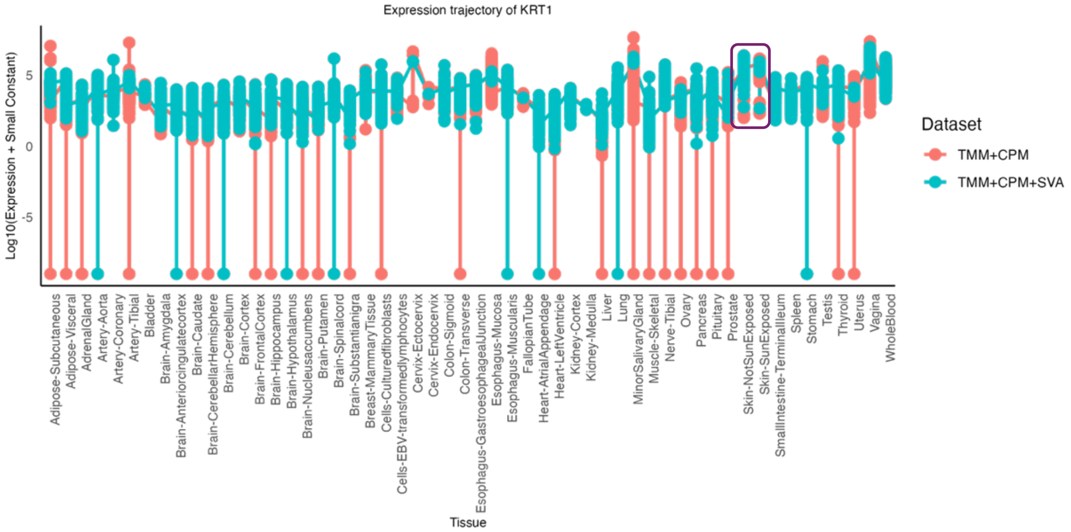


**Fig S2: ALB and KRT1 Expression Trajectories Across 54 GTEx Tissues**

(A) Trajectory plot of ALB expression across tissues (x-axis) with log-transformed values (y-axis). Purple rectangles highlight tissue-specific expression shifts between TMM+CPM and TMM+CPM+SVA processing. (B) Trajectory plot of KRT1 expression, with log-transformed values (y-axis). Purple rectangles indicate tissue-specific expression changes between processing methods.

Fig S3:

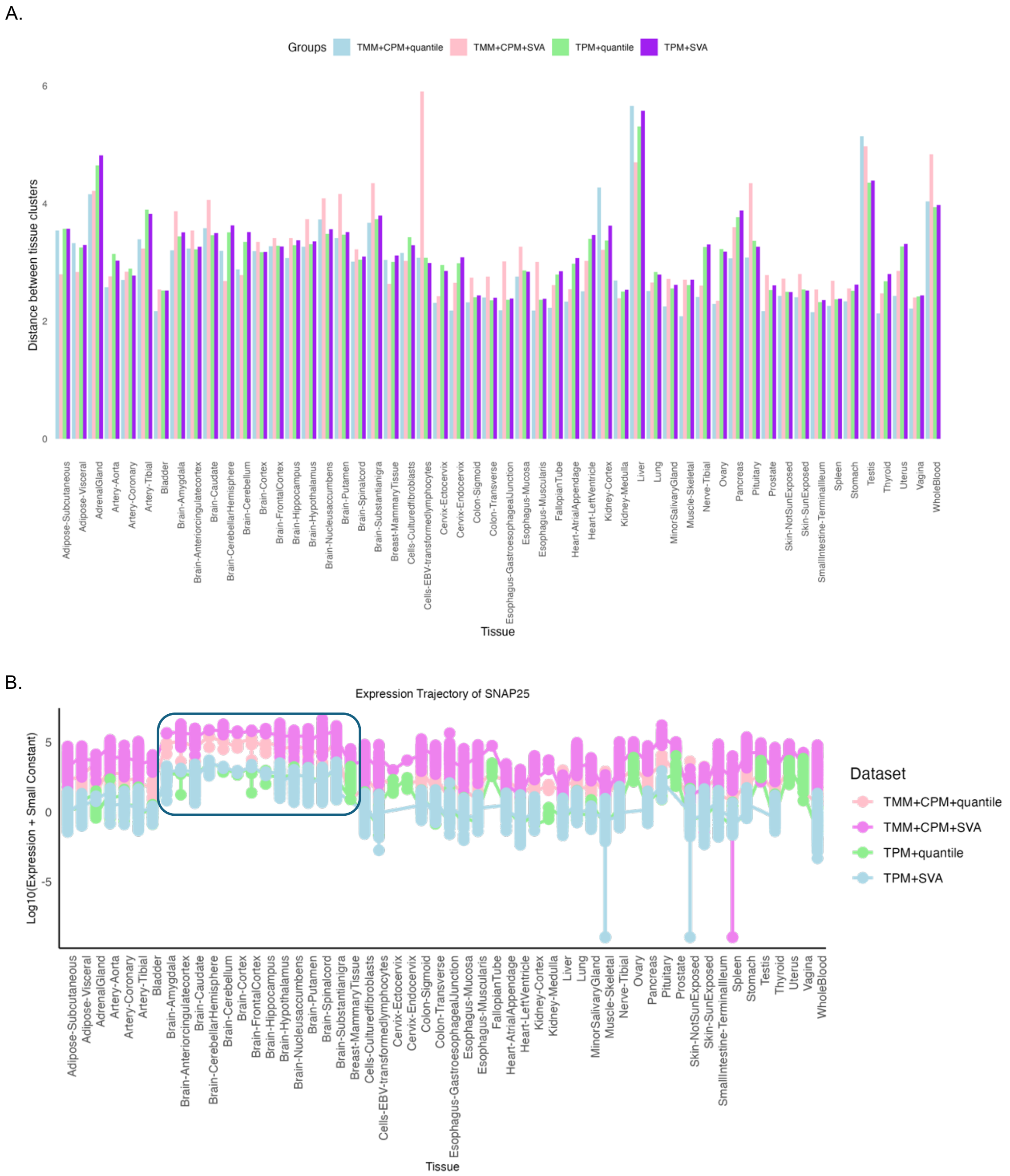


**Fig S3: Benchmarking processing methods across GTEx tissues.** A) Bar graph showing the distance between tissue clusters in GTEx gene expression data after preprocessing with different normalization pipelines: TMM+CPM+quantile (light blue), TMM+CPM+SVA (pink), TPM+quantile (light green), TPM+SVA (purple). B) Expression trajectory of SNAP25 across GTEx tissues for the TMM+CPM+quantile, TMM+CPM+SVA, TPM+quantile, and TPM+SVA processing pipelines, highlighting brain tissue-specific expression in purple.

Fig S4:

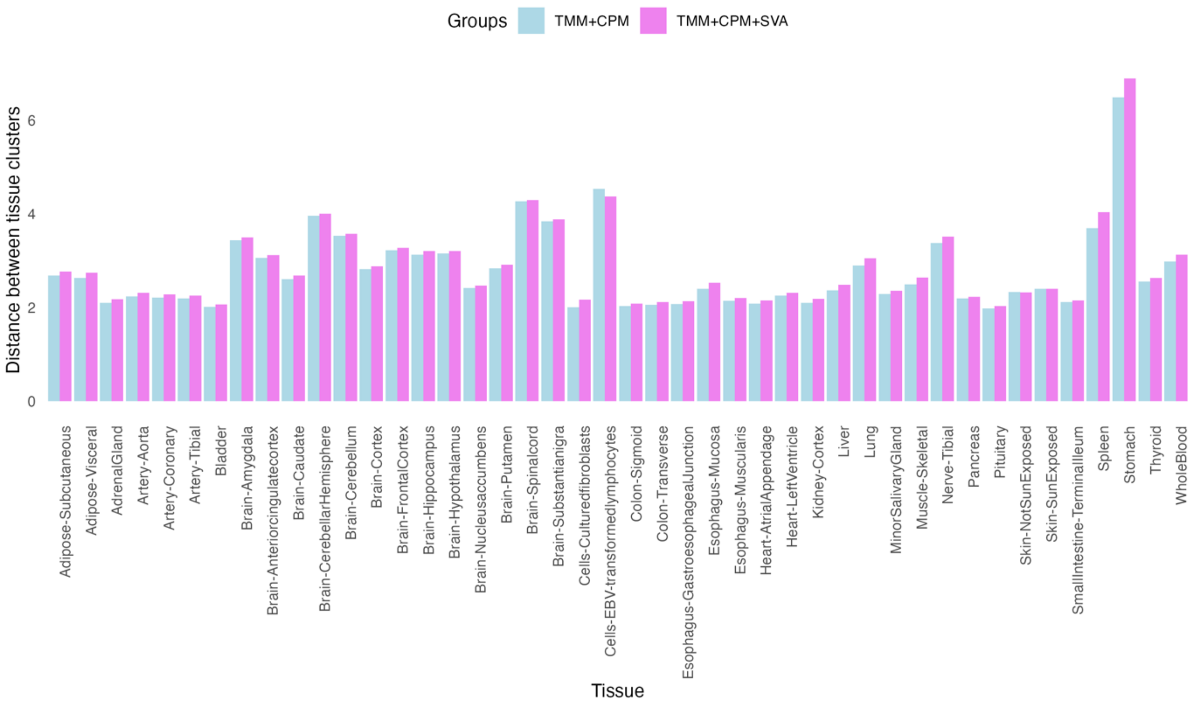


**Figure S4: Cross-validation of GTEx_Pro pipeline.**

Bar plot showing the average Euclidean distance between tissue clusters after TMM+CPM (light blue) and TMM+CPM+SVA (violet) processing, using an alternative gene set (AKT1, GSK3B, GDF11, FOXO1, SESN2, ULK1, PGC, PINK1, PDPK1, BCL2, HMOX1, FIS1, TNF, PARP1).

Fig S5:

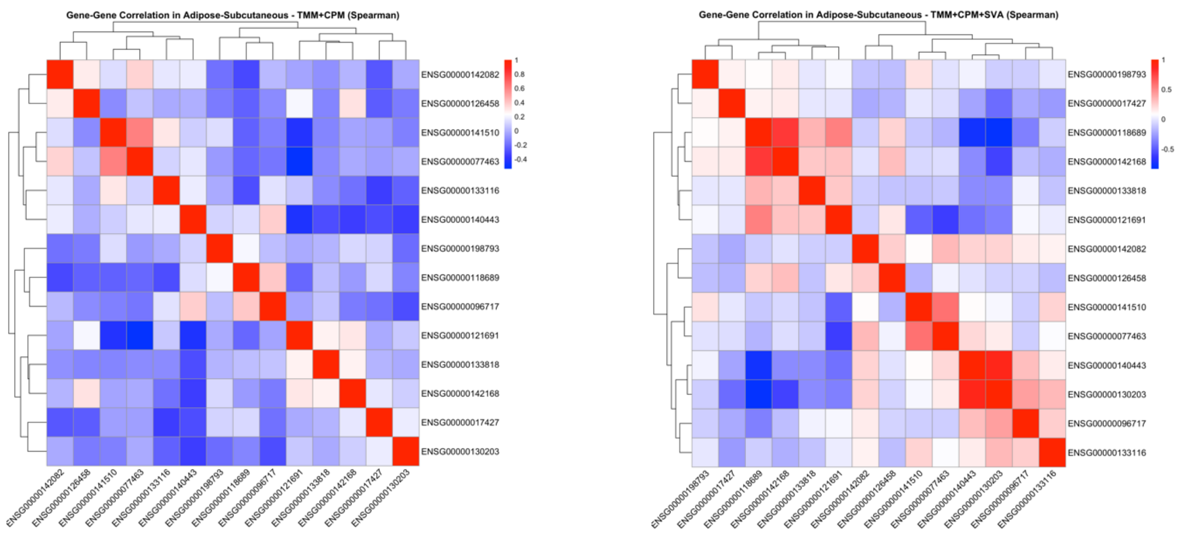


**Fig S5: Spearman correlation heatmap of gene expression in Adipose-Subcutaneous tissue.** Each cell represents the Spearman correlation coefficient (ρ) between genes, ranging from -1 (strong negative correlation, blue) to +1 (strong positive correlation, red). White indicates no correlation (ρ≈0). Heatmaps are shown for both TMM+CPM (before adjustment) and TMM+CPM+SVA (after adjustment), highlighting changes in gene-gene relationships due to SVA correction

Table 1: Sample count values for all the GTEx tissues generated in the normalization step.

sample_counts

| Tissue | SampleCount |
|---|---|
| Adipose-Subcutaneous.rds | 651 |
| Adipose-Visceral.rds | 541 |
| AdrenalGland.rds | 257 |
| Artery-Aorta.rds | 431 |
| Artery-Coronary.rds | 240 |
| Artery-Tibial.rds | 650 |
| Bladder.rds | 21 |
| Brain-Amygdala.rds | 152 |
| Brain-Anteriorcingulatecortex.rds | 176 |
| Brain-Caudate.rds | 246 |
| Brain-CerebellarHemisphere.rds | 215 |
| Brain-Cerebellum.rds | 241 |
| Brain-Cortex.rds | 255 |
| Brain-FrontalCortex.rds | 209 |
| Brain-Hippocampus.rds | 197 |
| Brain-Hypothalamus.rds | 202 |
| Brain-Nucleusaccumbens.rds | 246 |
| Brain-Putamen.rds | 205 |
| Brain-Spinalcord.rds | 159 |
| Brain-Substantianigra.rds | 139 |
| Breast-MammaryTissue.rds | 459 |
| Cells-Culturedfibroblasts.rds | 474 |
| Cells-EBV-transformedlymphocytes.rds | 165 |
| Cervix-Ectocervix.rds | 8 |
| Cervix-Endocervix.rds | 8 |
| Colon-Sigmoid.rds | 373 |
| Colon-Transverse.rds | 404 |
| Esophagus-GastroesophagealJunction.rds | 375 |
| Esophagus-Mucosa.rds | 553 |
| Esophagus-Muscularis.rds | 514 |
| FallopianTube.rds | 8 |
| Heart-AtrialAppendage.rds | 429 |
| Heart-LeftVentricle.rds | 430 |
| Kidney-Cortex.rds | 85 |
| Kidney-Medulla.rds | 4 |
| Liver.rds | 224 |
| Lung.rds | 576 |
| MinorSalivaryGland.rds | 162 |
| Muscle-Skeletal.rds | 790 |
| Nerve-Tibial.rds | 610 |
| Ovary.rds | 178 |
| Pancreas.rds | 327 |
| Pituitary.rds | 283 |
| Prostate.rds | 244 |
| Skin-NotSunExposed.rds | 604 |
| Skin-SunExposed.rds | 691 |
| SmallIntestine-TerminalIleum.rds | 187 |
| Spleen.rds | 240 |
| Stomach.rds | 357 |
| Testis.rds | 361 |
| Thyroid.rds | 651 |
| Uterus.rds | 141 |
| Vagina.rds | 154 |
| WholeBlood.rds | 733 |
